# Engineering Gac/Rsm signaling cascade for optogenetic induction of pathogenicity switch in *Pseudomonas aeruginosa*

**DOI:** 10.1101/2020.10.28.358515

**Authors:** Xinyi Cheng, Lu Pu, Shengwei Fu, Aiguo Xia, Shuqiang Huang, Lei Ni, Xiaochen Xing, Shuai Yang, Fan Jin

**Affiliations:** Hefei National Laboratory for Physical Sciences at the Microscale; Department of Polymer Science and Engineering, University of Science and Technology of China, Hefei 230026, China; CAS Key Laboratory of Quantitative Engineering Biology, Shenzhen Institute of Synthetic Biology, Shenzhen Institutes of Advanced Technology, Chinese Academy of Sciences, Shenzhen 518055, China

**Keywords:** Optogenetics, *Pseudomonas aeruginosa*, Gac/Rsm cascade, *Caenorhabditis elegans*, Pathogenicity control

## Abstract

Bacterial pathogens operate by tightly controlling the pathogenicity to facilitate invasion and survival in host. While small molecule inducers can be designed to modulate pathogenicity to perform studies of pathogen-host interaction, these approaches, due to the diffusion property of chemicals, may have unintended, or pleiotropic effects that can impose limitations on their use. By contrast, light provides superior spatial and temporal resolution. Here, using optogenetics we reengineered GacS of the opportunistic pathogen *Pseudomonas aeruginosa*, signal transduction protein of the global Gac/Rsm cascade which is of central importance for regulation of infection factors. The resultant protein YGS24 displayed significant light-dependent activity of GacS kinases in *Pseudomonas aeruginosa*. When introduced in *Caenorhabditis elegans* host systems, YGS24 stimulated the pathogenicity of PAO1 in BHI and of PA14 in SK medium progressively upon blue-light exposure. This optogenetic system provides an accessible way to spatiotemporally control bacterial pathogenicity in defined host even specific tissues to develop new pathogenesis systems, which may in turn expedite development of innovative therapeutics.

## Introduction

The conditions of the host, the pathogen, and the environment determine the result of a host-pathogen interaction ^1, 2^. Host-pathogen interactions are important to our understanding of the mechanisms of infectious disease, including its treatment and prevention. Besides, the location and timing of invasion or colonization and their subsequent mode of action in host decide the pathogen’s success of infection. However, there are few methods provide precise spatiotemporal control of infection. For instance, the injection systems applied to most host-pathogen interactions are not adequate for imaging the very early time course of infection due to the inevitable time delays of preparation of bringing pathogen in contacting with host cells. Other inoculation methods like feeding assay and immersion infection are also limited to control the occurrence of infection in defined location or tissue at specific times in intact systems ^3–5^. Accordingly, these techniques do not allow tracking the dynamics of host-pathogen interactions with spatiotemporal precision.

A promising approach to address the limitations is to couple the pathogenicity of bacteria to light signals. Light has advantages over chemical means of control of biological processes as it acts noninvasive, has low toxicity and most crucial, provides superior spatial and temporal resolution ^6^. To capitalize on these favorable properties, we, in this study, aimed to develop a light-controlled system by incorporating photoactive allosteric modulators semi-synthetically to enable previously unrivaled spatiotemporal control of pathogenicity. Given that the two-component system (TCS) GacS/GacA, is a signal transduction system that is of paramount importance for the regulation of infection behaviors and virulence factors in various gram-negative bacteria such as *Pseudomonas aeruginosa* (*P. aeruginosa*), a highly prevalent opportunistic human pathogen ^7^, we first sought to engineer a light-regulated histidine kinase optogenetic system based on GacS/GacA TCS in *P. aeruginosa* that could function in the involved Gac/Rsm signaling cascade and thus optogenetically control its pathogenicity. Here, we reprogrammed the input signal specificity of GacS by replacing its input sensor domain, which confer chemical sensitivity on its kinase activity, by the light-oxygen-voltage (LOV) blue light sensor domain of YtvA from *Bacillus subtilis*. Certain of the resulting fusion proteins retain kinase activity to activate the expression of sRNA, but are regulated by blue light instead of by chemical signal. We demonstrated and characterized one of the resultant fusion proteins named YGS24 that enabled the control of Gac/Rsm regulatory signaling cascade in a light-dependent manner with a low background activity and high light induction efficiency in *P. aeruginosa* (Figure 1A).

**Fig 1.**
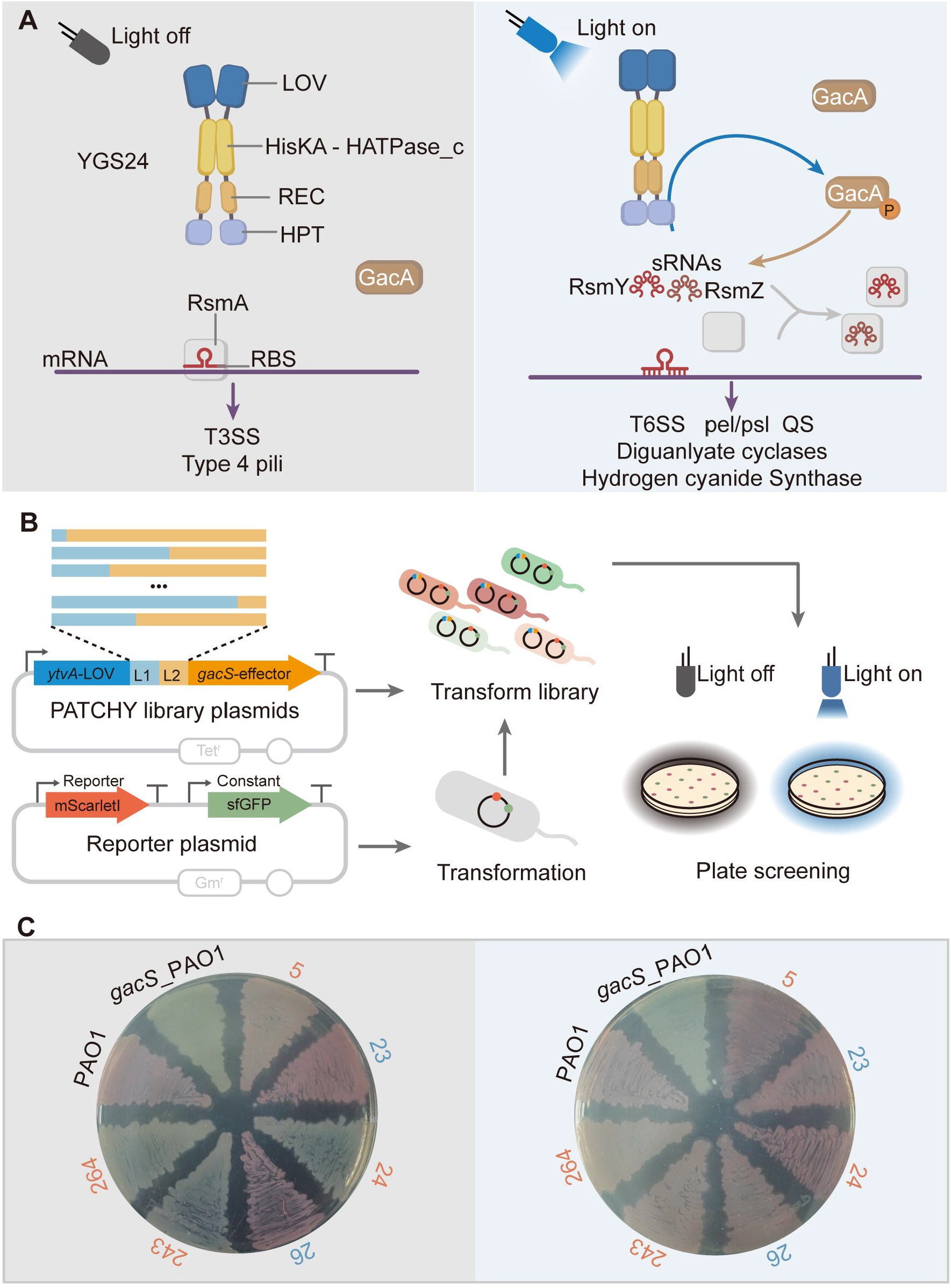
Development of the light-regulated YGSs. (A) Schematic representation of different states of the YGS/Rsm signaling cascade in light-off and light-on condition. (B) Construction and screening of the light-respondent YGSs. (C) Images of the LB plate assays of *rsmY* promoter response activity. Selected strains were grown on LB agar supplemented with the appropriate antibiotics, either under the illumination of darkness (left) or blue light (right). *P. aeruginosa* PAO1 wild type and *gacS*_PAO1 mutant with empty vector were here as control groups. Blue numbers, light-inactivated YGS proteins; orange numbers, light-activated YGS proteins.

We then tested the light-inducing pathogenicity switch of *P. aeruginosa* cells in *vivo* using PAO1-*Caenorhabditis elegans* (*C. elegans*) paralysis and PA14-*C. elegans* slow-killing model. In both pathogenesis systems, the irradiated engineered protein executed the inherent GacS histidine kinase activity involving the synthesis of cyanide and the expression of infection virulence factors respectively, hence resulting in the enhanced capability to kill *C. elegans*. Furthermore, we activated the Gac/Rsm signaling cascade of cells present in the lumen of *C. elegans* intestine by light and observed the lumen expansion due to colonization of PA14. Altogether, we regulated the infection signaling cascade of *P. aeruginosa* by an engineered light-responsive protein.

## Results

### Engineered fusion proteins YGSs displaying light-dependent kinase activity in *P. aeruginosa*

*B. subtilis* YtvA comprises an N-terminal LOV sensor domain adjacent to a C-terminal of Sulphate Transporter and AntiSigma factor antagonist (STAS) domain ^8^. The two domains are connected by a short α-helical linker ^9^. After activated by blue light, the dark-adapted state of YtvA-LOV domain (LOV450) reaches a blue-shifted covalent FMN-cysteine C4a-thiol adduct (LOV390) via the FMN triplet state (LOV660) within 2 μs ^10, 11^. In the absence of blue light LOV450 recovers within 2600 s ^12^. YtvA dimerizes in a head-to-head fashion. Jα helix relays the structural changes within the photosensor LOV module resulted from light absorption to alter the function of the effector STAS module ^13^.

GacS is the sensor kinase of a TCS that globally regulates the expression of acute and chronic virulence factors in *P. aeruginosa* ^14, 15^. The N-terminus of GacS contains two transmembrane regions. The cytoplasmic site of GacS contains an input domain Histidine kinases, Adenylyl cyclases, Methyl binding proteins, Phosphatases (HAMP) domain and three phosphotransfer domains: transmitter domain, receiver domain, and Hpt domain ^16^. The transmitter is made up of a HisKA domain and a HATPase-c domain. Upon receiving a stimulus through the input domain, the transmitter domain hydrolyses ATP to ADP using HATPase-c domain and auto-phosphorylates in the HisKA domain, then phosphorylates the receiver which phosphorylates the Hpt domain ^17^. The Hpt domain then transfers the phosphoryl group to the cognate response regulator GacA. Both GacS and GacA might also be active as dimers ^7^.

We produced the photoactivatable derivatives of GacS named YGSs by replacing the two transmembrane regions and the input domain of GacS with LOV domain of YtvA. To construct the photoreceptors YGSs, we applied the Primer-Aided Truncation for the Creation of Hybrid Proteins (PATCHY) method ^18^ for the efficient generation of hybrid gene library. Strains were generated by introducing each of YGSs, under the control of its native *gacS* promoter, into the *gacS* deletion mutant of PAO1, a wild type of *P. aeruginosa*. These strains were named a series of RO1-YGSs, for example of which is RO1-YGS24 that represents the strain carrying YGS24. And to facilitate the library screening and subsequent analysis, we engineered and exploited the transcription reporter of *rsmY* ^19^, one of the small RNAs that directly activated by the TCS of GacS/GacA, as a proxy for YGS/GacA/Rsm activity, in which an RNaseIII processing site located between the *rsmY* promoter and the RBS of *mScarletI* gene to ensure the independent translation of reporter gene, and introduced the reporter into RO1-YGSs cells. Accumulation expression of the fluorescent protein generates red colonies. We therefore screened the PATCHY libraries of YtvA-GacS fusions by color changes of colony appearance in the dark and light (Figure 1B). After winnowing about 384 colonies, six of these strains were found with obvious light-dependent *rsmY* reporter response activity, indicating the light-mediated kinase activity to regulate the expression of sRNA. We determined the nucleotide sequence of the corresponding YGSs by DNA sequencing (Figure S1). We showed that colonies of the cells that contain YGS23 or YGS26 faded upon blue light, reflecting light-inactivated kinase activity, and rest of the fusions, YGS5, YGS24, YGS243, and YGS264 displayed desired, light-activated kinase activity (Figure S1 and Figure 1C). We had a predilection for YGS24 due to its lower background activity and higher product of light-induced protein as judged by colony color and used it in all subsequent study (Figure 1C).

### Characterization of light-induced YGS24 activity in *P. aeruginosa*

As describe above, we focused on the YGS24. To further explore its light activated kinase activity in *P. aeruginosa*, we measured the activation of *mScarletI* transcriptional reporter for *rsmY* at single-cell level as a ratio to a constitutively expressed *sfgfp* reporter in the cells of RO1-YGS24. We verified that the increase in the fluorescence ratio of mScarletI-to-sfGFP (RFP-to-GFP ratio, RG ratio) was mainly a result of the increased activity of the *rsmY* promoter as opposed to the little changes in the sfGFP normalization control. Single cells have the significant induced *rsmY* expression within 2 h under constant 470 nm light of 96.4 μW/cm^2^ (Figure 2A and B). The mean fluorescence intensity of cell populations of RO1-YGS24 in darkness kept at low level as well as the *gacS* mutant, while the fluorescence level of cells under light reached comparable with that of wild type PAO1 (Figure 2C and Figure S2). To determine the light intensity dependence of YGS24 activity, we measured the RG ratio for single cells in culture medium after illumination for 6 h with 470 nm of varying intensity. Even a light intensity of around 1 μW/cm^2^ can partially activate, and an intensity of approximately 25 μW/cm^2^ can completely activate the blue light dependent *rsmY* expression (Figure 2D). Moreover, the YGS24-based optogenetical system was demonstrated to exhibit fully reversible activation of the Gac/Rsm signaling cascade (Figure 2E). Additionally, we noticed that the cells of RO1-YGS24 grown under darkness showed slightly faster growth than the cells exposed to light, and a similar growth defect was observed in wild type cells comparable to the cells of *gacS* mutant, which indicated that growth inhibition was correlated with the activation of the Gac/Rsm pathway, excluding the possibility that phototoxic effects are responsible for this phenomenon (Figure S3).

**Fig 2.**
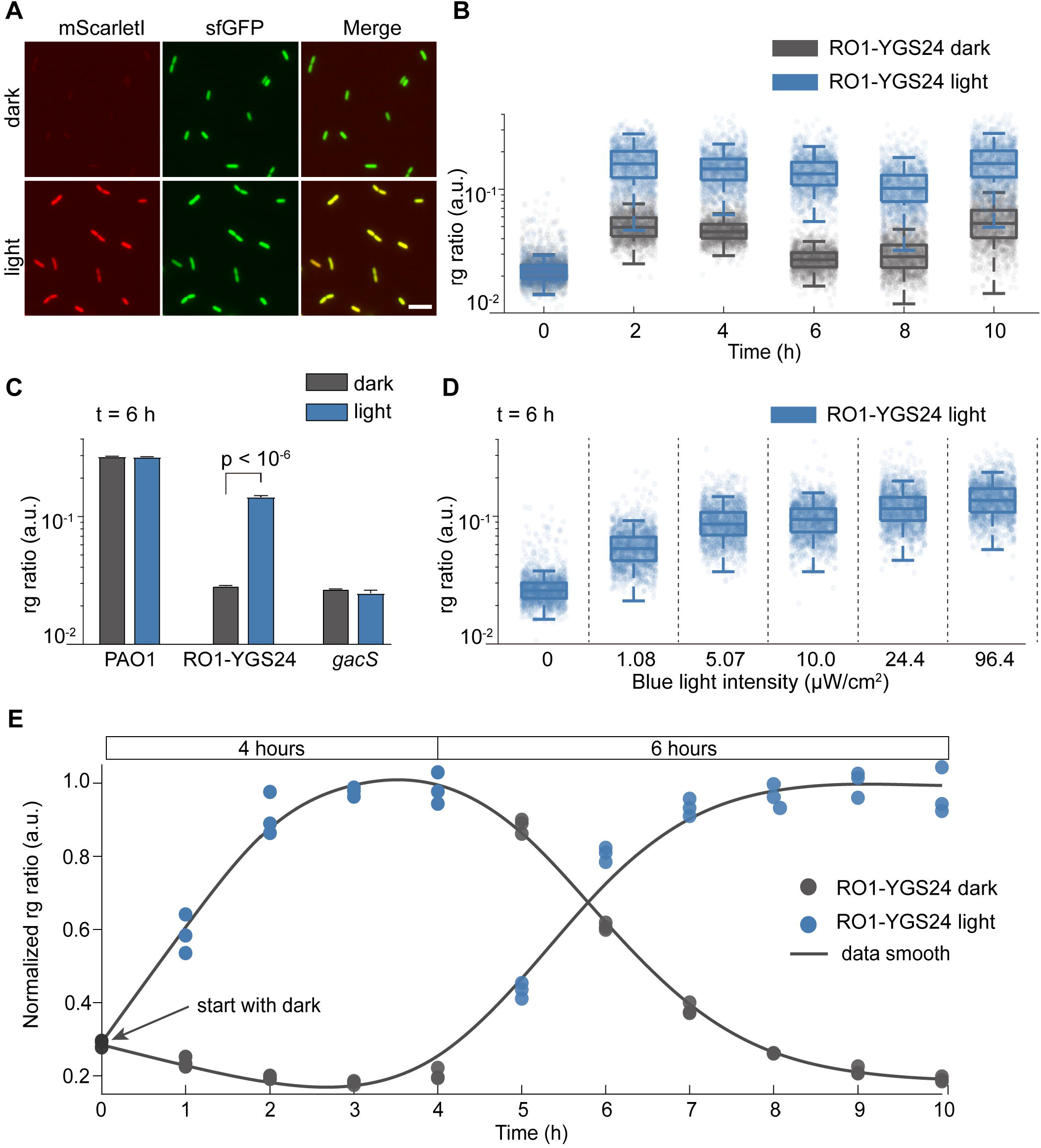
Performance of light-activated protein YGS24 in *gacS*_PAO1 mutant. Strains were transformed with a *rsmY* reporter encoding mScarletI fluorescent protein. A *sfgfp* reporter fused to constitutive promoter J23102 served as an internal control. (A) The mScarletI, sfGFP, and corresponding merged fluorescence microscopy images of RO1-YGS24 under blue light (bottom) and darkness (up). Scale bar, 5 μm. (B) Boxplot of rg ratio separated by cells illuminated with and without blue light. Every dot in the plot diagram represents one counted cell; more than 10^3^ cells from 3 replicates were analyzed. (C) Rg ratio of wild type PAO1, RO1-YGS24, and *gacS*_PAO1 measured after 6 h under blue light or darkness. Data represent mean values of 10^3^ single-cell events; error bars represent 1□SD of n□=□3 biologically independent samples. (D) Rg ratio of RO1-YGS24 measured after blue light illumination of different intensities for 6 h. (E) Rg ratio measured at indicated times during incubation with illumination under continuous blue light for 4 h and then by 6 h darkness, or with inverse illumination condition. Every circle represents mean values of 10^3^ single-cell events; data are representative of n□=□3 independent experiments.

### Induction of pathogenicity switch of cells through blue light in *P. aeruginosa*-*C. elegans* models

The Gac/Rsm pathway globally controls the expression of virulence factors in *P. aeruginosa.* For investigating the pathogenicity of *P. aeruginosa*, the nematode *C. elegans* has been chosen as an alternative host.

It has been shown that *P. aeruginosa* PAO1 kills *C. elegans* within 4 h when the strain is grown on rich brain-heart infusion (BHI) medium and *gacS* mutant is strongly attenuated in this virulence model ^20, 21^. To identify the virulence switch of RO1-YGS24 cells in the conditions of light and dark, we performed a paralytic killing assay with *C. elegans* as the pathogenesis system. In this assay, cells growing under constant 470 nm light of 120 μW/cm^2^ killed up to 100% of the nematodes after 4 h, while only about 20% of total worms killed by cells growing in darkness (Figure 3A).

**Fig 3.**
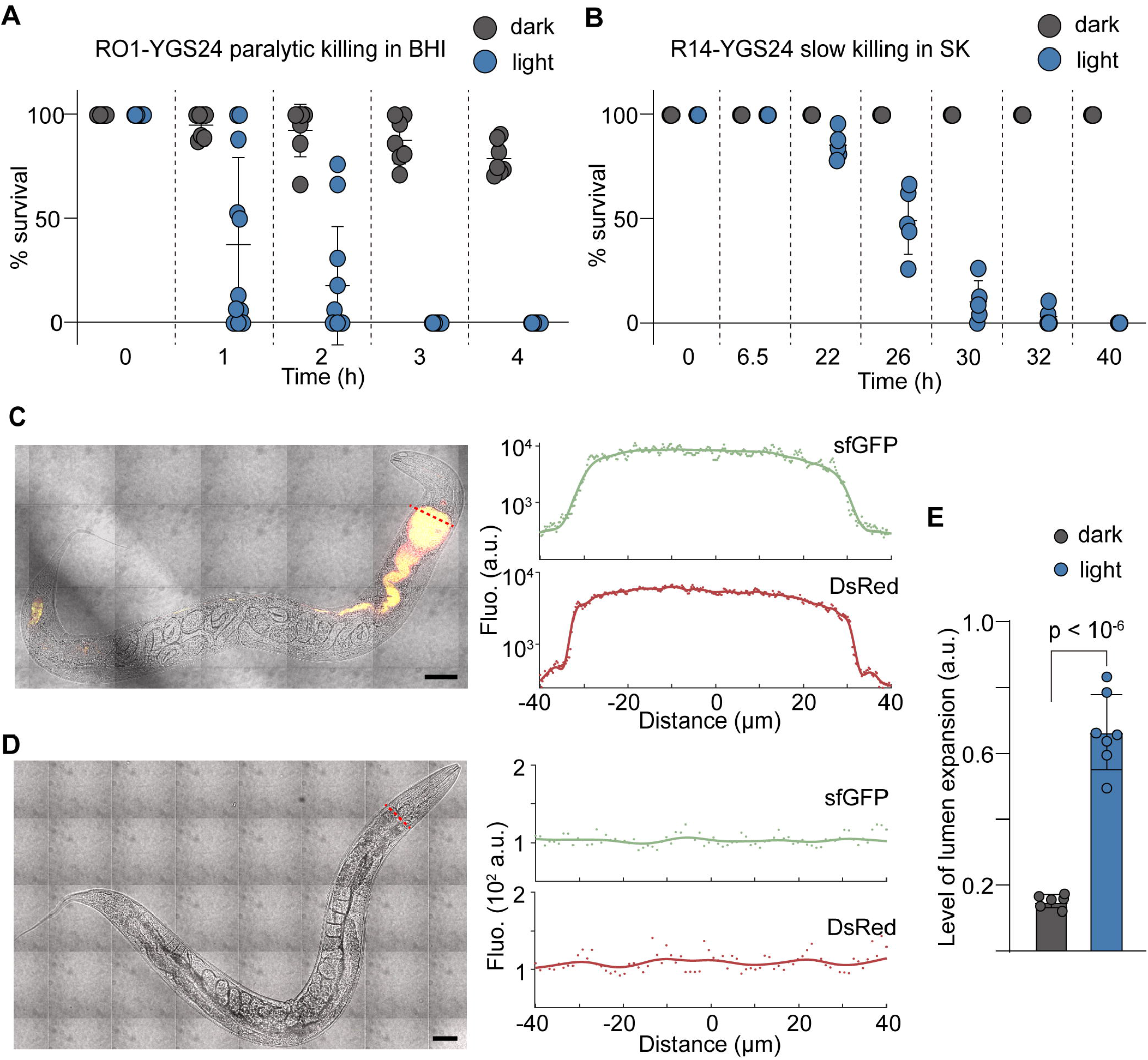
*P. aeruginosa-C. elegans* killing assays. (A) *C. elegans* survival assays comparison between the strain RO1-YGS24 or (B) R14-YGS24 with and without blue light. Every circle represents data of one biologically independent experiment. (C) The entire lumen of worms fed on illuminated cells of R14-YGS24-(RP2-sfGFP-DsRed.T3) for 20 h was filled with fluorescence-expressing cells. In contrast, (D) the lumen of worms fed on cells growing under darkness for same time showed little detectable fluorescence. Scale bar, 50 μm. (E) The degree of pharynx expansion of worms fed on R14-YGS24-(RP2-sfGFP-DsRed.T3) under continuous blue light was significantly higher than that of worms fed on cells under darkness. Data represent the mean ± SD from 7 biologically independent samples; every circle represents individual data point from one worm.

For the other hand, when *C. elegans* is fed on *P. aeruginosa* PA14 grown on SK agar plate, the mechanism by which *P. aeruginosa* kills *C. elegans* involves an infection-like process that correlates with the accumulation of cells in the lumen of *C. elegans* intestine ^22^. The killing occurs over the course of several days and is referred to as slow killing. Mutation in the *gacS* gene of PA14 was severely defective in the ability to kill *C. elegans* under slow-killing conditions ^23^. We then constructed the strain named R14-YGS24, using similar strategy with RO1-YGS24 to replace the *gacS* naturally present in the wild type PA14 with the gene of YGS24, and examined whether light functions on our engineered cells in this killing model. We fed *C. elegans* on R14-YGS24 cells growing under the conditions of illumination and darkness respectively. As the detailed statistics shown in Figure 3B, the worms died within 20-40 h under illumination conditions, while no killing was observed during the experiment period in darkness, indicating that the unilluminated cells are nonpathogenic. We further confirmed that the cells that have the deletion of *gacS*, either in PAO1 or PA14, are seriously deficient in killing *C. elegans*, which are similar with the RO1-YGS24 or R14-YGS24 cells under darkness, while the lethality of the RO1-YGS24 or R14-YGS24 cells on *C. elegans* under light conditions are comparable with that of wild type cells (Figure S4). Taken together, these results showed that we had the ability to optogenetically regulate the virulence of *P. aeruginosa* cells ranging from the level of *gacS* mutant to wild type with the use of engineered photo-responsive protein YGS24.

To verify that light induced the signal pathways of *P. aeruginosa* we had already known in engineered cells, we designed translational fusion reporter of *hcnB* in RO1-YGS24 and of *clpV1* in R14-YGS24, *hcnB* associates with cyanide poisoning and *clpV1* is connected with infection virulence, as a proxy for the pathogenicity in its corresponding killing model ^24, 25^ (for details see Methods). We showed that the illumination induction strongly boosted the expression of the cyanide poisoning marker *hcnB* in RO1-YGS24 and the chronic virulence marker *clpV1* in R14-YGS24 (Figure S5), in accordance with the results of killing assays.

To further monitor the living state of *C. elegans* during the infection process, we engineered the R14-YGS24 cells expressing sfGFP and DsRed.T3 to feed nematodes, in which DsRed.T3 was constitutively expressed for visualization of cells and sfGFP was used as the proxy for sRNA expression. After 20 h of feeding on illuminated cells, as shown in Figure 3E, both green and red fluorescence was observed throughout the expanded lumen of worm intestines, indicating the accumulation of cells with activated Gac/Rsm cascade. In contrast, there are few cells present, indicated by very little fluorescence, in the lumen of worms fed on unilluminated cells at this time point (Figure 3C and D). The level of expansion of lumen of worms, which we roughly defined as the ratio of widths of the pharynx section of lumen to the body part of *C. elegans* at measured time, stayed below 0.2 for the unilluminated ones, while it increased to 0.68±0.1 for the illuminated ones after 20 h feeding with R14-YGS24. (Figure 3E). Collectively, these results suggested that the light induced activation of Gac/Rsm cascade of PA14 cells had the ability to accumulate in the worm gut accompanied by lumen expansion gradually, thereby resulting in the lethal infection.

### Regulation of *P. aeruginosa* Gac/Rsm cascade in *C. elegans* with spatial resolution

One specific advantage of light-activated systems over other induced system is the superior degree of spatial control, so we sought to activate the infection cascade of bacterial cells in the intestine of *C. elegans*. We performed it using PA14-*C. elegans* slow-killing model mentioned above to ensure a sufficient induction time before worm died. Given that the unilluminated engineered cells are nonpathogenic and barely replicate in the worm intestine, we mixed wild type cells to help the engineered cells to colonize in the worm gut. We fed *C. elegans* on the mixed cells under darkness for 25 h and transferred the worms to a microfluidic device ^26^ (Figure 4A and Figure S6) for spatial light induction. We monitored the cells with intermittent microcopy imaging. After illumination for 2 h, green fluorescence was observed within the illuminated worms, indicating the induced expression of sRNA and the expression switch of downstream genes of Gac/Rsm pathway (Figure 4B and C). Following illumination to 10 h, green fluorescence increased more than 3-fold (Figure 4B and C). In contrast, the green fluorescence barely changed in unilluminated worms (Figure S7). At this time point, we noticed that the worms, both under darkness and illumination condition, had an expansion of lumen and became immobile, and this might be mainly caused by the continued virulence of wild type cells from mixture feeding.

**Fig 4.**
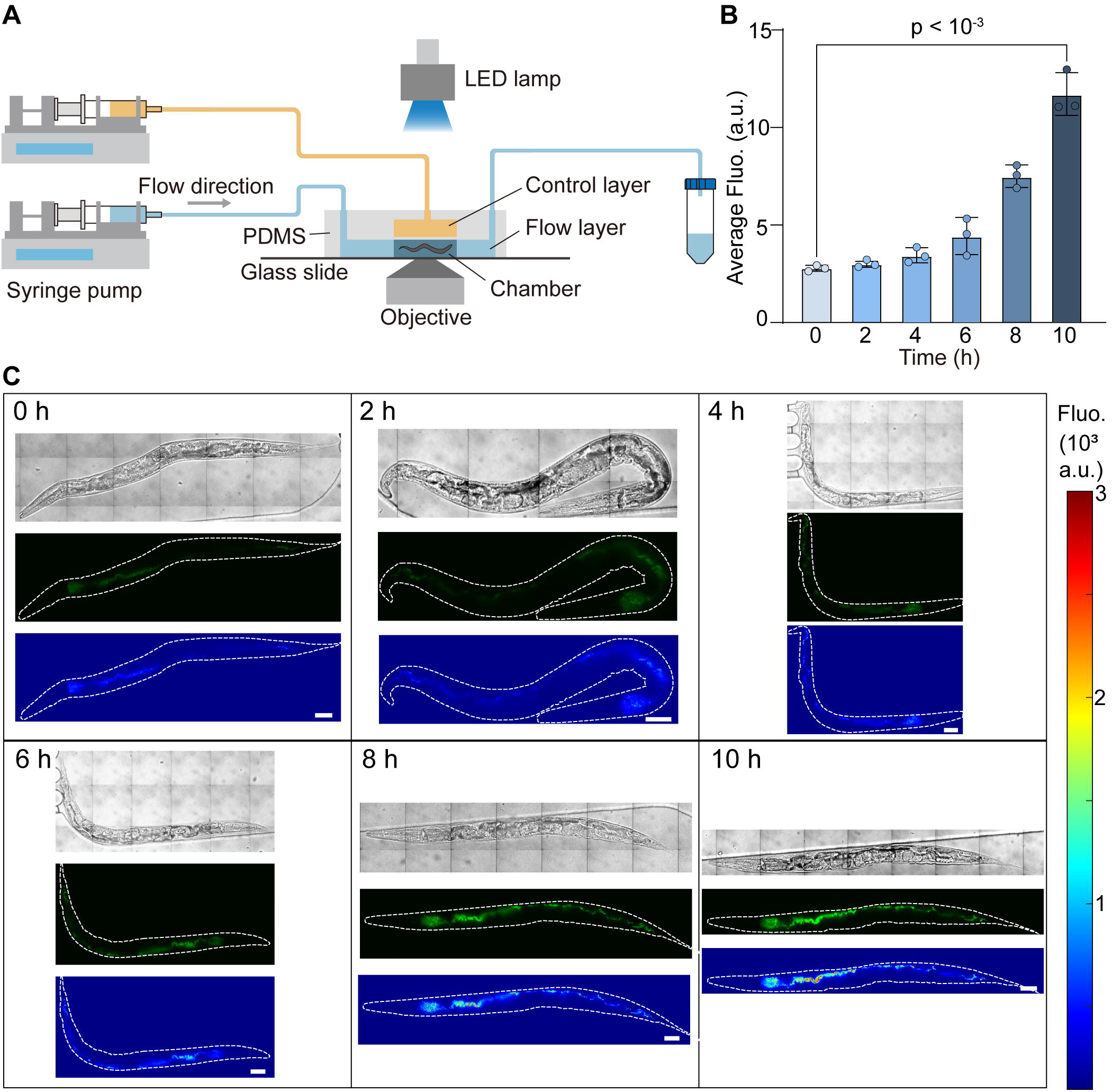
Light-induced infection process in *P. aeruginosa-C. elegans* model. (A) Schematic representation of experiment for activating YGS/Rsm signalling cascade of *P. aeruginosa* in *C. elegans* in the microfluidic device by blue light. (B) Cells accumulated over time in the lumen of *C. elegans* illuminated with blue light. Data represent the mean ± SD from□3 biologically independent samples; every circle represents individual data point from cells in one worm. (C) Bright-field (upper), sfGFP fluorescence (middle) microscopy images and heat map for fluorescence intensity (lower) showed spatial control over blue-light-induced gene expression from R14_YGS24-(RP2-sfGFP-DsRed.T3) in the intestine of *C. elegans*. Used as control group some worms fed on same condition and transferred to dark environment (S7 Fig). Scale bar, 50 μm.

## Discussion

Light offers a more precise, efficient and reversible way to manipulate biological functions directly at the protein level. In the present study, we constructed the photoreceptor YGSs by recombining the blue light responsive LOV photosensor module of *B. subtilis* YtvA with the *P. aeruginosa* GacS histidine kinase effector (Figure 1A and Figure S1). Optogenetic protein interaction switches employ conformational changes of specific proteins, such as LOV domain proteins, to control protein−protein interactions by light ^27^. However, the applications of optogenetics have mainly been in eukaryotic cells ^28^. And these photo-responsive proteins designed in bacteria were mainly in model strain such as *Escherichia coli* to enable spatiotemporally transcriptional activation of user-defined exogenous or endogenous genes from specific promoter under light illumination and could be further used to manipulate many biological processes ^29–31^. Nevertheless, adoption of the transgene systems to other species of bacteria may introduce interference with endogenous proteins or genes due to homology ^32, 33^. These light-switchable gene expression systems, most importantly, are incapable of regulating global regulatory cascades partly because of large numbers of downstream gene, which severely restricts the utilization of such optogenetic systems for further systemic research that involves complex genetic regulation such as pathogenicity.

The resultant protein YGS24 showed light-dependent kinase activity of GacS, thus allowing us to reversibly stimulate the global Gac/Rsm signaling cascade which is of central importance for the regulation of infection and virulence factors in *P. aeruginosa* by blue light (Figure 1C and Figure 2). However, poor penetration of blue light through nontransparent tissues should be considered for the further use in research of cell cultures or animal models. The insights emerged from YGSs engineering were expected to facilitate engineering of new light-activated optogenetic tools fusing a photosensory module encompassed the spectral region of needed and a sensor kinase of TCSs.

When tested in host models, our engineered optogenetic system enabled the light-regulated pathogenicity switch of bacterial cells and correspondingly tunes the susceptibility of *C. elegans* to *P. aeruginosa*-mediated killing (Figure 3). In the two *P. aeruginosa*-*C. elegans* models, unilluminated strains RO1-YGS24 and R14-YGS24 were incapable to kill *C. elegans*, namely, light induced innocuous-to-mortal virulence switch of pathogen. We expected to control analogous morbidity switch in various environment or host, as the Gac/Rsm cascade in *P. aeruginosa* and other gram-negative bacteria regulated different phases of pathogenicity. The absence of GacS of clinical *P. aeruginosa* discomposed the Gac/Rsm pathway leading to depletion of the sRNAs RsmY/RsmZ and, in consequence, to expression of type 3 secretion system (T3SS) or type 4 pili, hallmark of acute infection, while switching off the expression of virulence factors promoting chronic infection such as T6SS, hydrogen cyanide synthase or Pel polysaccharides ^15^ (Figure 1A). This suggested that we might control acute-to-chronic virulence switch in *P. aeruginosa* by blue light.

*P. aeruginosa* PA14 kills *C. elegans* involving a chronic infection-like process that associates with the colonization and accumulation of cells in the lumen of *C. elegans* intestine in SK medium ^22^. Due to hardly adherence of strain R14-YGS24-(RP2-sfGFP-DsRed.T3) in the lumen of *C. elegans* intestine under dark condition, we fed worms on commingled cells of R14-YGS24-(RP2-sfGFP-DsRed.T3) and PA14 and induced the innocuous-to-mortal virulence switch of *P. aeruginosa* by blue light. And because of restless of worms, they were temporally immobilized by the microfluid device for less than one minute, and lacking in efficient tracking approach, we regulated spatially the accumulation of cells within single worm, instead of in the part of lumen of one worm or at single-cell revolution theoretically.

Host-pathogen interactions can be investigated on many levels, considering that not all interactions lead to disease and those that do have a complex progression that heads to this state. Moreover, the genetic uniqueness and diversity of each host or pathogen and the inherent variability in the complexity and types of host-pathogen interactions result in an immense and somehow unfathomable array of different combinations. Each individual host-pathogen interaction relationship is unique, and the pathogen would show different attributes of aggressive and toxicity to various tissues even within one host ^34^. Overall, we engineered a new light-activated protein to optically control the infection and virulence factors in *P. aeruginosa*, thereby further expanding the tool set in the area of pathogenic research that has great demands for precise spatiotemporal control of bacterial pathogenicity.

## Materials and methods

### Microbiological methods

The plasmids, strains and primers used in this study are listed in Table S1-S3 in Supporting Information respectively. Standard molecular cloning techniques were used for construction of related plasmids in *E. coli* strain Top10. Deletion mutants *gacS*_PAO1 and *gacS*_PA14 were constructed by using well-established protocols for *P. aeruginosa* based on two-step allelic exchange ^35^. To screen the library and test the performance of YGS24, we exploited the transcriptional reporter of *rsmY*, named RP1-mScarletI-sfGFP. To construct this reporter plasmid, PCR fragments, promoter P*rsmY*, an RNaseIII processing site, red fluorescent protein *mScarletI* and terminators (T0T1) were assembled into one piece by overlap extension PCR ^36^. Another fragment J23102-*sfgfp*-T0T1 was aligned by the same way. After the overlap PCR, the two resulting aligned fragments were then cloned into pJN105 through multiple cloning sites. The *hcnB* translational reporter HP-mScarletI-sfGFP and *clpV1* translational reporter CP-mScarletI-sfGFP were constructed by replacing P*rsmY*-RNaseIII with translational promoter P*hcnB* and P*clpV1* respectively. To visualize bacteria more clearly in the intestine of *C. elegans*, we construct another reporter RP2-sfGFP-DsRed.T3 by replace *mScarletI* and *sfgfp* with *sfgfp* and *dsRed.T3* respectively using PCR and Gibson assembly ^37^. All PCR reactions here were performed with a high-fidelity polymerase (Phanta Super-Fidelity DNA Polymerase, Vazyme). Unless performing *C. elegans* assays, both *E. coli* and *P. aeruginosa* stains were grown in LB medium or on LB agar. Plasmids were maintained in media supplemented with the following concentrations: gentamicin (Aladdin) 15 μg/mL (pJN105, pUCP20Gm and derivatives), tetracycline (Aladdin)10 μg /mL (pUCP20Tet and derivatives) in *E. coli*; gentamicin 30 μg/mL (pJN105, pUCP20Gm and derivatives), tetracycline 100 μg/mL (pUCP20Tet and derivatives) in *P. aeruginosa*. *C. elegans* N2 strain was kindly shared by the lab of Shouhong Guang (University of Science and Technology of China), and was grown on *E. coli* (OP50) lawns on nematode growth media agar (NGM) at 20°C. To obtain a synchronized population, gravid adult nematodes were transferred by wire pick to fresh NGM plates with OP50 for 2–4□hours to lay eggs, and then removed.

Cells of RP1-mScarletI-sfGFP were used for YGS24 light-sensitive assay. Strains were streaked on LB agar from frozen stocks and incubated overnight at 37°C. Single colony was inoculated a 1ml culture of LB broth in the dark at 30°C until OD600 reached approximately 2.4. The culture was diluted to an OD600 of approximately 0.05 and incubated under blue light or in the dark at 30°C /250 rpm. Fluorescence intensities were measured every 2 hours. Growth curves were measured under the same condition giving an initial diluted OD600 of ~0.02. The translational reporter HP-mScarletI-sfGFP and CP-mScarletI-sfGFP were transformed into RO1-YGS24 and R14-YGS24 respectively. The two strains were cultured overnight followed by 1:100 dilution to 1 mL LB medium and cultured for 14 h (RO1-YGS24-(HP-mScarletI-sfGFP)) or 11 h (R14-YGS24-(CP-mScarletI-sfGFP)) at 30°C /250 rpm under blue light or darkness, then fluorescence intensities were measured. For the intensity curves measurement, light condition was changed from blue light to dark, or on the contrary. Overnight strain culture was diluted to an OD600 of ~0.05 and then incubated in shaker with 250 rpm at 30°C in the dark until OD600 reached approximately 0.6. The culture was then diluted to an OD600 of ~0.3 and incubated in the shaker under certain light conditions. Every hour, the strain culture was diluted to an OD600 of ~0.3 and the fluorescence intensity was measured.

For PATCHY library screening, Petri dishes were placed under 3 W/m^2^ blue light emitting from an LED lamp (λ= 470 nm, M470L2, Thorlabs) or in the dark. For *C. elegans* assays, blue light intensity from LED was set at approximately 1.2 W/m^2^.

### PATCHY library

PATCHY library was generated by using a modified version of the published protocol^18^. The *Bs*YtvA-LOV domain and the three phosphotransfer domains from *Pa*GacS with their full-length linkers were cloned in tandem in the pUCP20. This constructing plasmid was functioned as the template. The forward and reserve primers annealed to different positions within the C-terminal LOV module linker and the linker of the N-terminal phosphotransfer domains respectively and were each shifted in their annealing sequence by one triplet to avoid frame shift. A one-pot PCR reaction with two sets of forward and reserve primers was carried out to amplify the template. To reduce the template in PCR products, we reduced the dosage of the template to 0.002 ng/μL in the PCR reaction. The PCR products were purified by PCR-clean up (AxyPrep PCR Cleanup Kit, Axygen) and then self-circularized via T4 Polynucleotide Kinase (ThermoFisher), ATP (ThermoFisher) and T4 DNA Ligase (ThermoFisher). The reaction mix was transformed into chemically competent *E. coli* Top10. The plasmids were extracted and transform into PAO1 with pJN105-P*rsmY* reporter (RP1-mScarletI-sfGFP). Cells harboring the construct and the reporter were grown in LB medium with tetracycline (100 μg/mL) and gentamicin (30 μg/mL) for 20 h at 37°C. Single colonies were then inoculated 96-deep-well plates containing 600 μL LB (with tetracycline and gentamicin) per well for 20 h at 37°C. 2 μL of each single colony solution were grown on LB medium (with tetracycline and gentamicin) at 30°C under blue light and in the dark to screen effective fusions.

### Paralytic killing assay

*P. aeruginosa* strains were incubated in shaker with 250 rpm at 37°C overnight in Brain heart infusion (BHI) (Oxoid) broth and then diluted 100-fold into fresh broth. BHI agar plates (60 mm diameter) were spread with 200 μL each of the dilution and then incubated for 24 h at 37°C to form bacteria lawns. Gentamicin was added at 30 μg/mL if required. Adult nematodes from stock plates were collected in a minimal volume of M9 buffer (22□mM KH_2_PO_4_, 42□mM Na_2_HPO_4_, 86□mM NaCl, 1□mM MgSO_4_). Droplets containing nematodes were spotted onto the *P. aeruginosa* lawns. The plates were then sealed with parafilm and incubated at room temperature. The paralysis of the nematodes was scored with a microscope. The assays of RO1-YGS24 strain with gentamicin resistance were always under blue light or in the dark during the whole experiments. Each plate was scored at only one time point, we prepared same plates for different time point at each experiment.

### Slow killing assay

The slow killing assay was performed by using a modified protocol of previously described methods ^22, 38^. 50 μL overnight cultures of *P. aeruginosa* stains (wild-type PA14, *gacS*_PA14, R14-YGS24) were pipetted onto 60 mm diameter modified SK agar plates (tryptone instead of peptone was used) with or without 50 μg/mL tetracycline, and were spread by using sterile L spreaders to entirely cover the plates. The plates were incubated at 37°C for 24 h and then grown at 25°C for an additional 24 h. Synchronized adult worms were seeded on plates. The killing plates were incubated at 25°C and scored for live worms at intervals. The assays of R14-YGS24 were performed under blue light and dark conditions, respectively.

### Microfluidic experiment

The microfluidic chip was designed to enable individual worm loading, long-term culture, harmless immobilization and fluorescence imaging. The chip was fabricated in PDMS (poly(dimethylsiloxane)), which is gas permeable, transparent, non-fluorescent and biocompatible with organisms. The chip was composed of three layers: top control layer, middle PDMS membrane and bottom flow layer (Figure S5). The bottom flow layer consisted of worm inlet, loading channels and eight chambers (2 mm length, 500 μm width, and ~60 μm height) for worm culture and imaging. The captured worm was prevented from escaping due to the narrow grooves (50 μm width, 20 μm interval distance) at the end of chamber. The deformable PDMS membrane (40 μm thick) was assembled between the bottom flow layer and top control layer. By activating the microvalve in the control layer, the PDMS membrane deformed to the chamber in the flow layer and worms could be immobilized.

To observe bacteria behavior *in vivo*, a cell mixture comprised of 40 μL overnight culture of PA14 harboring pJN105 and pUCP20Tet empty vectors and 10 μL overnight culture of R14-YGS24 carrying RP2-sfGFP-DsRed.T3 was cultured in darkness to prepare bacteria lawn. Synchronized young-adult worms were fed for 25 h in darkness and washed with M9 buffer to wipe bacteria off. Fresh M9 buffer contained with worms was added into the inlets in the flow channel of microfluidic device. After worms swam into the loading channel, they were loaded into the culture chambers under the M9 flow driven by syringe pump, and the device was placed in blue light. For imaging, worms were immobilized by controlling the microvalve in the control layer. To immobilize worms, the valve was switched on by positive pressure of water from the control layer. Then the deformable PDMS membrane was pushed up, to squeeze the worms to the side of the imaging channels so that worms could be immobilized and imaged. After imaging, valve was switched off by removing pressure and then the worms could be released and recover their mobile activity quickly.

### Microscopy

In light-sensitive experiments, fluorescent images were collected by using an invert fluorescent microscope (Olympus, IX71) equipped with a 100× oil immersion objective and two sCMOS cameras (Zyla 4.2, Andor). The GFP (sfGFP) and RFP (mScarletI) were excited by using 488/10 nm and 567/15 nm lasers respectively and the emission filters used were 520/28 nm and 631/36 nm respectively. For the fluorescent intensity curves measurement, fluorescence imaging was performed using a spinning-disk confocal (CSU-X1; Yokogawa) inverted microscope (IX81; Olympus) equipped with a laser combiner system (Andor Technology), a 100× oil immersion objective (Olympus) and EMCCD camera (iXon897). To collect the fluorescence of sfGFP, a 488 nm laser was used as exciter, a 524/40 nm channel was used as emitter. The mScarletI was excited using a 561 nm laser and the fluorescence was detected by a 605/40 nm channel.

In *C. elegans* imaging assays, bacteria lawns were prepared by the same method in slow killing assays. Age-synchronized young-adult worms were collected for each experiment. Nematodes were fed with illuminated and unilluminated bacteria lawns of R14-YGS24 carrying RP2-sfGFP-DsRed.T3 for 20 h in light and darkness respectively. To observe bacteria behavior in vivo, mixture contained with 40 μL overnight culture of PA14 carrying pJN105 and pUCP20Tet and 10 μL overnight culture of R14-YGS24 carrying RP2-sfGFP-DsRed.T3 was cultured in darkness to prepare bacteria lawn. Worms were fed for 25 h in darkness and washed with M9 buffer to wipe bacteria off. M9 buffer contained with worms was added into EP tube or microfluidic device and placed in blue light or darkness for 10 h. Images were collected using spinning disk confocal inverted microscope as mentioned above, a 60× oil immersion objective and an EMCCD camera (iXon 897). The sfGFP or DsRed.T3 fluorescence was excited using a 488 or 561□nm laser respectively and imaged with two different emission channels (524 or 605□nm).

### Statistical analysis

Images acquired were analyzed using ImageJ software or MATLAB codes ^39^. Cell masks were obtained using images of fluorescent protein which driven by J23102, and then the reporter FP fluorescence was measured by computing the mean intensity within masks. To obtain mean fluorescence intensity, we subtracted the average fluorescence intensity per pixel of the background from the average intensity per pixel in the given cell. Mean values and standard deviations were obtained from at least three independent experiments (biological replicates). Statistical analysis was performed with Student’s t-test using GraphPad Prism version 8.0.

## Supporting Information

**Table S1.** A list of plasmids used in this study; **Table S2.** A list of strains and nematode used in this study; **Table S3.** A list of primers used in this study **Figure S1.** Alignment of linker regions of light-regulated YtvA-GacS fusions from PATCHY libraries; **Figure S2.** The mean fluorescence intensity of cell populations under blue light; **Figure S3.** Activation of the GacS leads to reduced growth; **Figure S4.** P. aeruginosa-C. elegans nematode model; **Figure S5.** Light induced corresponding signal pathways of P. aeruginosa in engineered cells; **Figure S6.** Schematic of the microfluidic device for the feeding, illumination, fixing and observation of worms; **Figure S7.** Microscope images of worms fed on mixture cells

## Author Information

### Author Contributions

X.C., L.P., S.Y. and F.J. designed the project; X.C., L.P. and S.F. carried out the experiments and participated in data analysis; S.H. designed and provided the microfluidic devices; A.X., L.N. and X.X. contributed methodological of microscopy; S.Y. performed the majority of data analysis; and X.C., L.P. and S.Y. wrote the manuscript.

## Acknowledgements

We thank professor Shouhong Guang of university of science and technology of China for gifting us C. elegans N2. This work was supported by National Key Research and Development Program of China (Grant no. 2020YFA0906900 and Grant no. 2018YFA0902700 to F.J.) and National Natural Science Foundation of China (Grant no. 31901028 to S.Y., Grant no. 21774117 to F.J., Grant no. 31700745 to X.X. and Grant no. 31700087 to L.N.) and China Postdoctoral Science Foundation (Grant no. 2020M672881 to S.Y.). The funders had no role in study design, data collection and analysis, decision to publish, or preparation of the manuscript.

