## Supplementary figures and images for "Engineering Gac/Rsm signaling cascade for optogenetic induction of pathogenicity switch in *Pseudomonas aeruginosa*"

### Figure S1

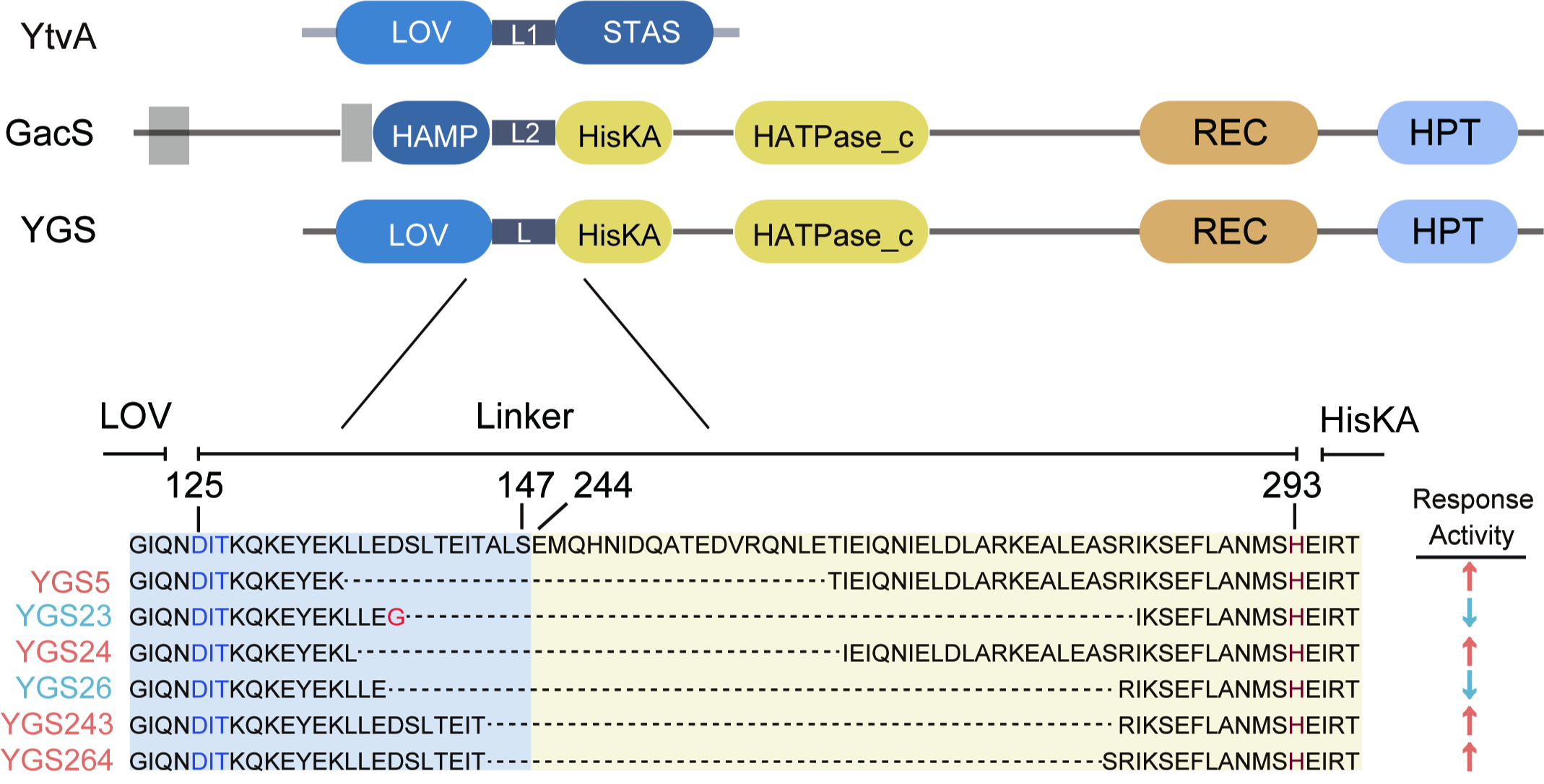

### Figure S2

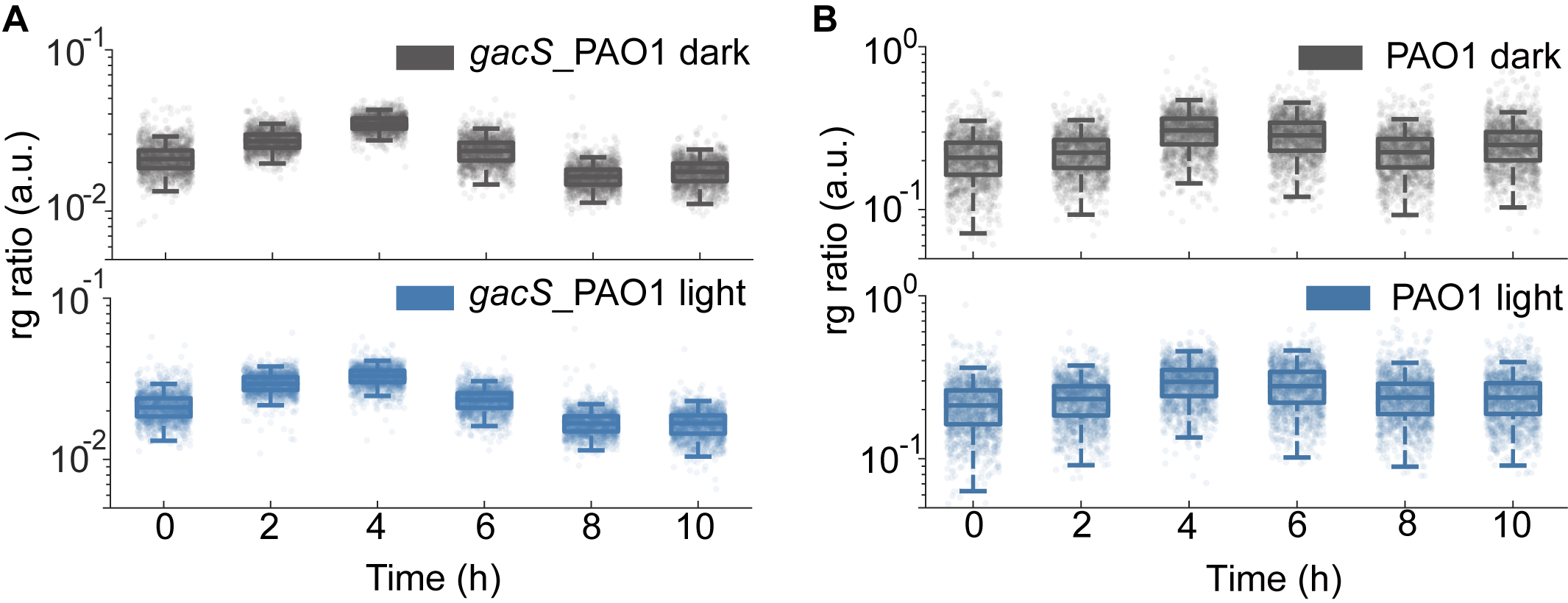

### Figure S3

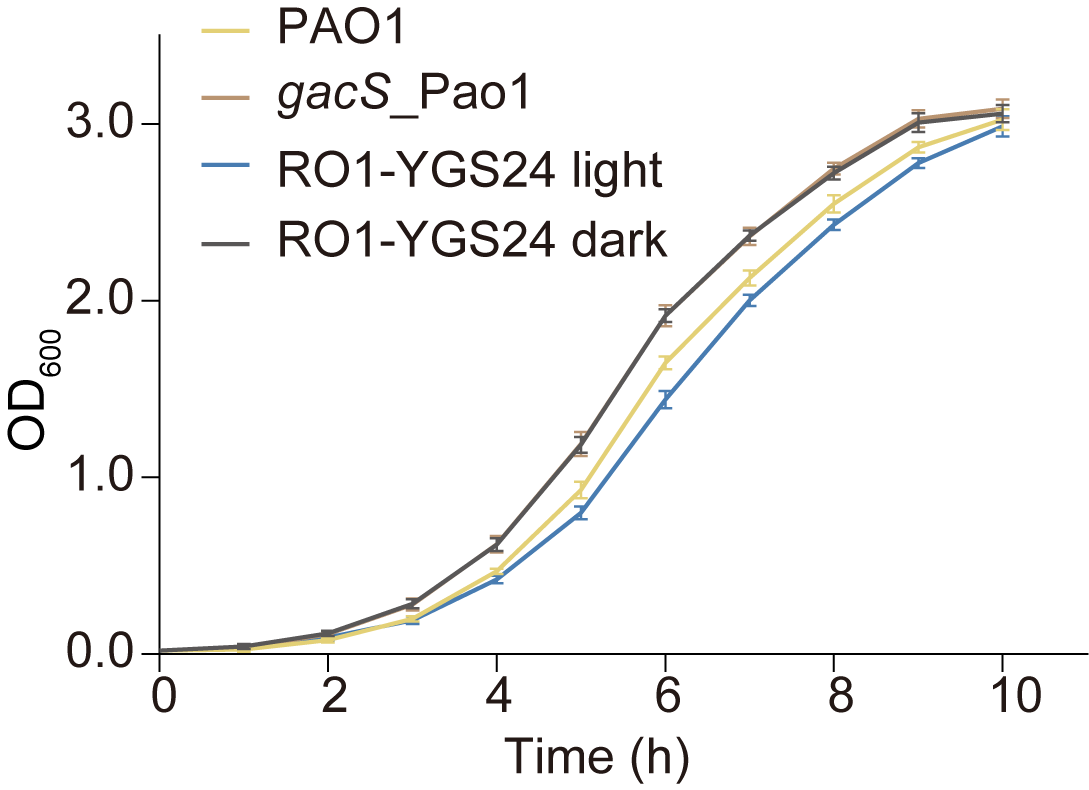

### Figure S4

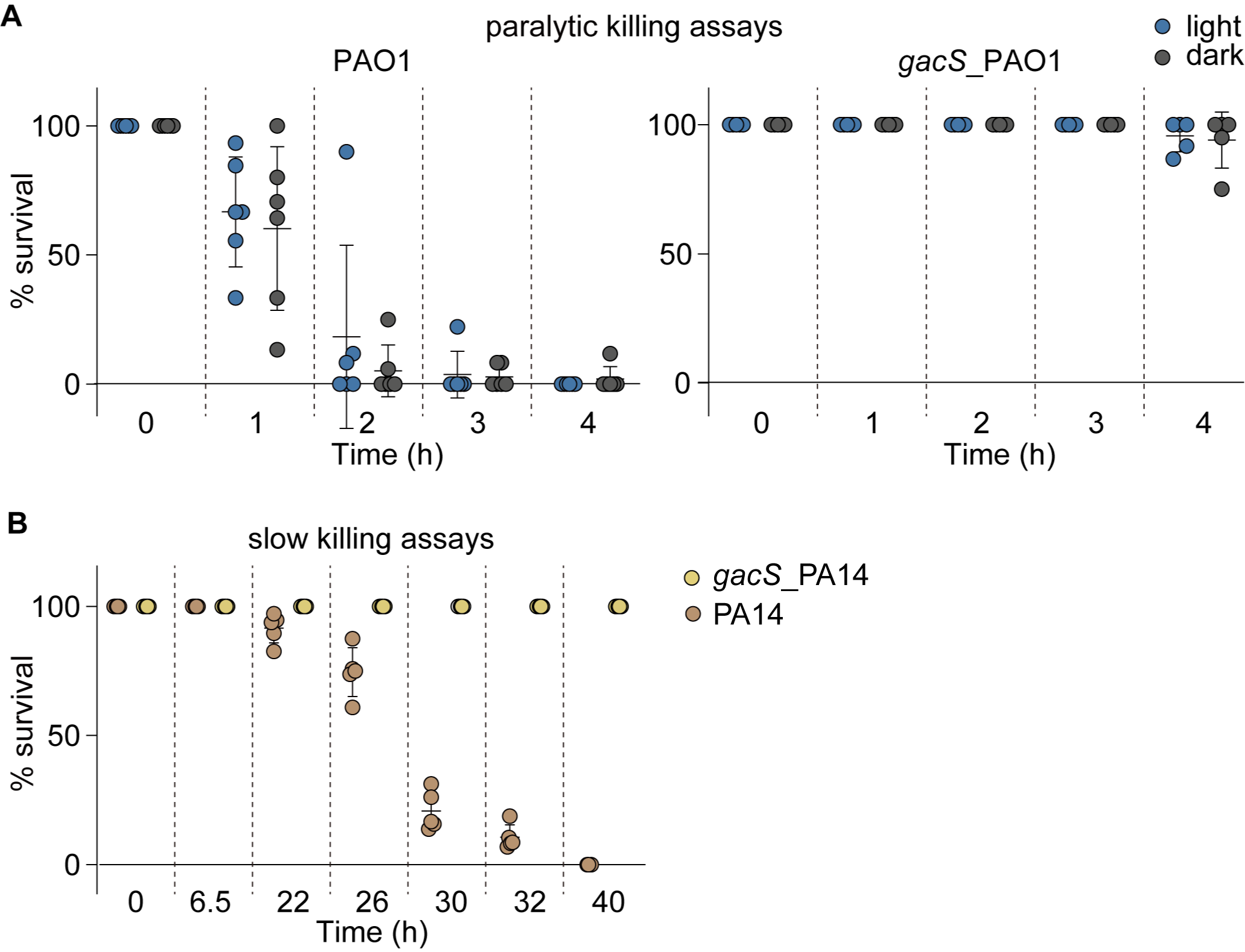

### Figure S5

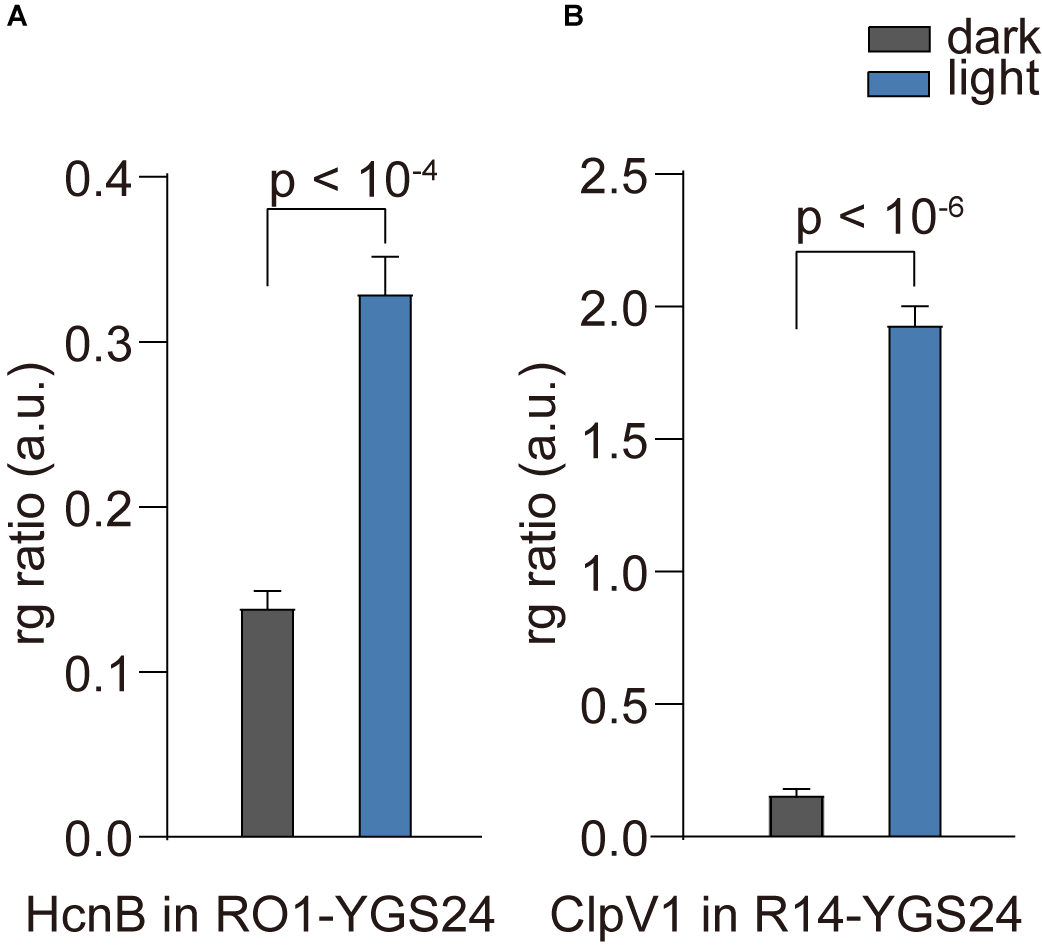

### Figure S6

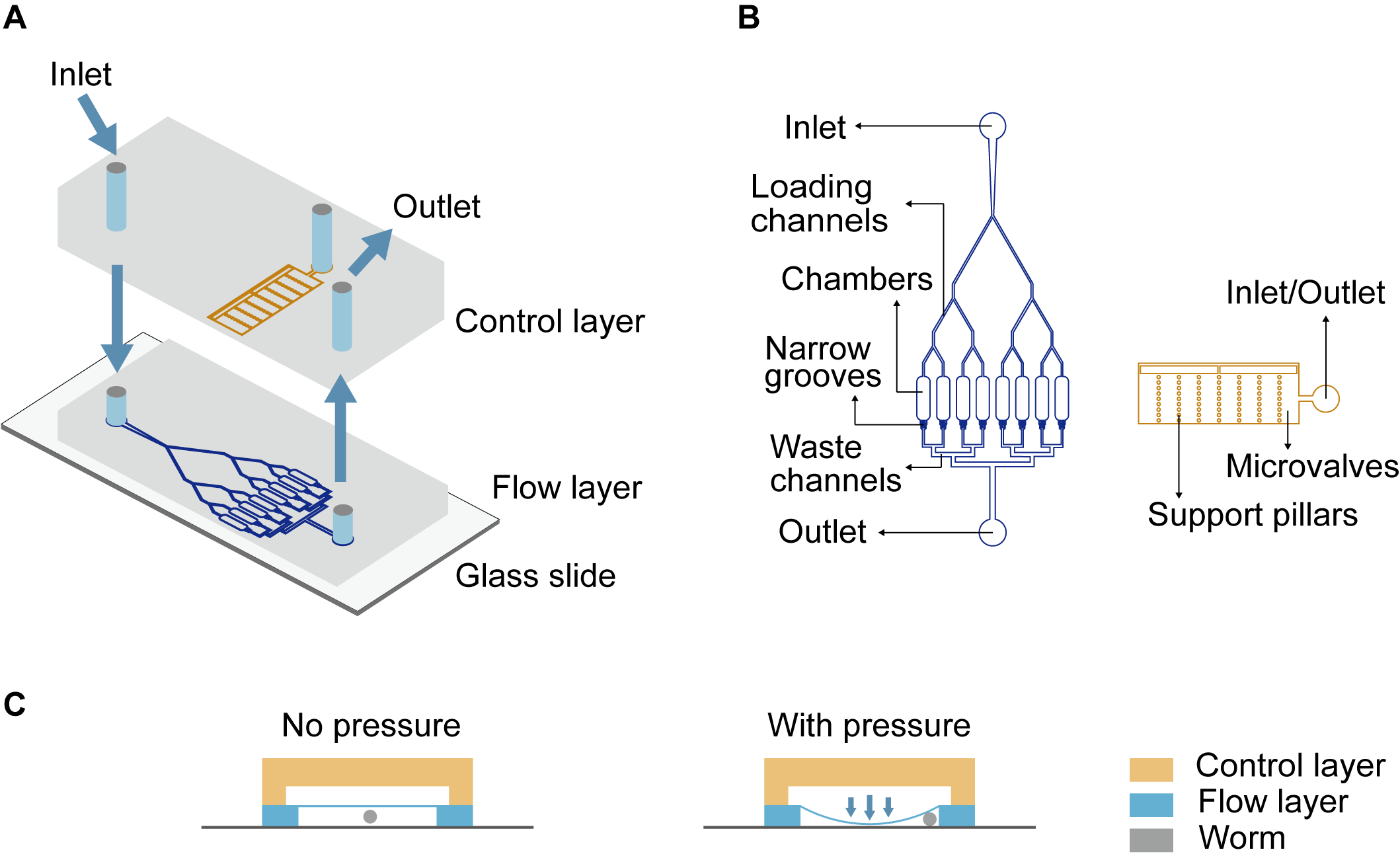

### Figure S7

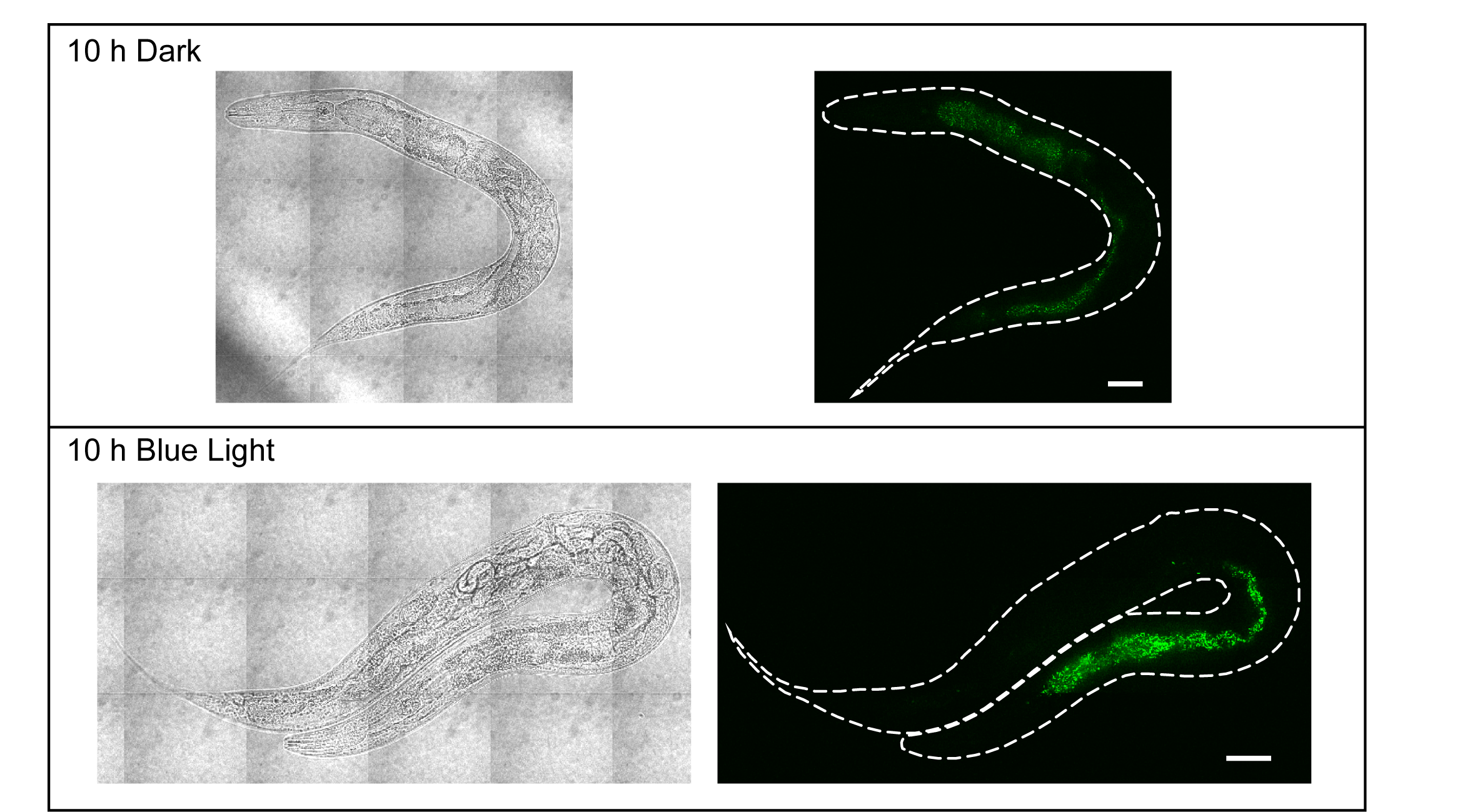
