## Supplementary material for "Engineering Gac/Rsm signaling cascade for optogenetic induction of pathogenicity switch in *Pseudomonas aeruginosa*": Table S1

| Plasmids | Description | Reference/source |
|---|---|---|
| pEX18Gm | Allelic-exchange vector with MCS from pUC18; oriT sacB lacZ; $Gm^{r*}$ | H. P. Schweizer |
| pEX18Gm_*gacS* | pEX18Gm-derived allelic-exchange vector for *gacS*; $Gm^r$ | This study |
| pUCP20Tet | Ampicillin resistance cassette of pUCP20 replaced with tetracycline resistance fragment; $Tet^r$ | This study |
| pUCP20Gm | Ampicillin resistance cassette of pUCP20 replaced with gentamicin resistance fragment; $Gm^r$ | This study |
| RP1-mScarletI-sfGFP | RsmY transcriptional reporter *mScarletI* driven by the *rsmY* promoter cloned into pJN105; $Gm^r$ $P_{rsmY}$-*RNaseIII*-*mScarletI*-$T_0T_1$-J23102-B0034-*sfgfp*-$T_0T_1$-pJN105 | This study |
| RP2-sfGFP-DsRed.T3 | RsmY transcriptional reporter *sfgfp* driven by the *rsmY* promoter cloned into pJN105; $Gm^r$ $P_{rsmY}$-*RNaseIII*-RBS2-*sfgfp*-$T_0T_1$-J23102-B0034-*dsRed.T3*-$T_0T_1$-pJN105 | This study |
| pUCP20-YGS | The *Bs*YtvA-LOV domain and the three phosphotransfer domains from *Pa*GacS with their full-length linkers cloned in tandem in pUCP20Tet; $Tet^r$ | This study |
| pUCP20Gm-YGS24 | *ygs24* driven by the *gacS* promoter cloned into pUCP20Gm; $Gm^r$ | This study |
| pUCP20Tet-YGS24 | *ygs24* driven by the gacS promoter cloned into pUCP20Tet; $Tet^r$ | This study |
| HP-mScarletI-sfGFP | HcnB translational reporter *mScarletI* driven by the translational *hcnB* promoter cloned into pJN105; $Gm^r$ $P_{hcnB}$-*mScarletI*-$T_0T_1$-J23102-B0034-*sfgfp*-$T_0T_1$-pJN105 | This study |
| CP-mScarletI-sfGFP | ClpV1 translational reporter *mScarletI* driven by the translational *clpV1* promoter cloned into pJN105; $Gm^r$ $P_{clpV1}$-*mScarletI*-$T_0T_1$-J23102-B0034-*sfgfp*-$T_0T_1$-pJN105 | This study |
| ⁎ $Gm^r$, gentamicin resistance; $Tet^r$, tetracycline resistance. | | |
