## Supplementary material for "Engineering Gac/Rsm signaling cascade for optogenetic induction of pathogenicity switch in *Pseudomonas aeruginosa*": Table S2

| Bacterial strains | Description | Source |
|---|---|---|
| *P. aeruginosa* | | |
| PAO1 | Wild-type strain | J.D.shrout |
| PAO1-(pUCP20Gm) | PAO1 strain harboring pUCP20Gm; Gm$^{r*}$ | This study |
| PAO1-(pUCP20Tet)-(RP1-mScarletI-sfGFP) | PAO1 strain harboring RP1-mScarletI-sfGFP and pUCP20Tet; Gm$^{r}$ and Tet$^{r}$ | This study |
| *gacS*_PAO1 | Deletion of *gacS* in PAO1; nonresistant | This study |
| RO1-(pUCP20Gm) | *gacS*_PAO1 strain harboring pUCP20Gm; Gm$^{r}$ | This study |
| RO1-YGS24 | *gacS*_PAO1 strain harboring pUCP20Gm-YGS24; Gm$^{r}$ | This study |
| RO1-(RP1-mScarletI-sfGFP) | *gacS*_PAO1 strain harboring RP1-mScarletI-sfGFP; Gm$^{r}$ | This study |
| RO1-(pUCP20Tet)-(RP1-mScarletI-sfGFP) | *gacS*_PAO1 strain harboring RP1-mScarletI-sfGFP and pUCP20Tet; Gm$^{r}$ and Tet$^{r}$ | This study |
| RO1-YGS24-(RP1-mScarletI-sfGFP) | *gacS*_PAO1 strain harboring RP1-mScarletI-sfGFP and pUCP20Tet-YGS24; Gm$^{r}$ and Tet$^{r}$ | This study |
| RO1-YGS24-(HP-mScarletI-sfGFP) | *gacS*_PAO1 strain harboring HP-mScarletI-sfGFP and pUCP20Tet-YGS24; Gm$^{r}$ and Tet$^{r}$ | This study |
| PA14 | Wild-type strain | J.D.shrout |
| PA14-(pJN105) - (pUCP20Tet) | PA14 strain harboring pJN105 and pUCP20Tet; Gm$^{r}$ and Tet$^{r}$ | This study |
| *gacS*_PA14 | Deletion of *gacS* in PA14; nonresistant | This study |
| R14-YGS24 | *gacS*_PA14 strain harboring pUCP20Tet-YGS24; Tet$^{r}$ | This study |
| R14-YGS24-(RP2-sfGFP-DsRed.T3) | *gacS*_PA14 strain harboring RP2-sfGFP-DsRed.T3 and pUCP20Tet-YGS24; Gm$^{r}$ and Tet$^{r}$ | This study |
| R14-YGS24-(CP-mScarletI-sfGFP) | *gacS*_PA14 strain harboring CP-mScarletI-sfGFP and pUCP20Tet-YGS24; Gm$^{r}$ and Tet$^{r}$ | This study |
| *E. coli* | | |
| OP50 | Uracil auxotroph. | Shouhong Guang |
| Top10 | F-*mcrA* Δ(*mrr-hsRMS-mcrBC*)Φ80*lacZ*Δ*M15* Δ*lacX74 recA1 ara*D139 Δ(*araleu*)7697 *galU galK rpsL*(Nal$^{R}$) endA1 *nupG* | Invitrogen |
| Top10- (pUCP20-YGS) | Top10 strain harboring pUCP20-YGS; Tet$^{r}$ | This study |
| Nematode | | |
| *C. elegans* | Wild-type N2 strain | Shouhong Guang |
| * Gm$^{r}$, gentamicin resistance; Tet$^{r}$, tetracycline resistance. | | |
