## Supplementary material for "Engineering Gac/Rsm signaling cascade for optogenetic induction of pathogenicity switch in *Pseudomonas aeruginosa*": Table S3

| Primer purpose and name | DNA sequence |
| --- | --- |
| Deletion of *gacS* | |
| Up-GacS-F | AAGGGGGATGTGCTGCAAGGCGATTAAATCGTGTTCAGCCATGGAG |
| Up-GacS-R | CGCCGAACAACTGAGCGAAACCCTGCTCAAG |
| Dw-GacS-F | GCAGGGTTTCGCTCAGTTGTTCGGCGATCAGTTG |
| Dw-GacS-R | GGGATCCTCTAGAGTCGACCTGCAGCAGCCATACCGATAGCAGGA |
| PATCHY library Construction | |
| Pgacs-GacS-F | CATGCCTGCAGGTCGACTCTAGAGGGGCAGCCAAATAAGATTGATTAGCAA |
| Pgacs-GacS-R | AACAGGAGTCCAAGCTCAGCTAATTGGTCAGAGTTCGCTGGAG |
| GacS-HisKA-F | ACAACATCGACCAGGCCACCGAGGACGTACGGCAGAACC |
| PgacS-R | ACGTCTCTCCGTCGAGC |
| YtvA-LOV-F | CCGATCTGGCTCGACGGAGAGACGTGTGGCTAGTTTTCAATCATTTGGGAT |
| YtvA-LOV-R1 | TGAAAGTGCAGTAATTTCCGTGAGGGAATCCTCGAGAAGCTTTTCATATTCTTTTTGCTTGGTGATATCATTCTGAATTCCGACAAAATAC |
| YtvA-LOV-R2 | GGTGGCCTGGTCGATGTTGTGCTGCATTTCTGAAAGTGCAGTAATTTCCGTGAG |
| Patchy-F1 | TGAAAGTGCAGTAATTTCCGTGAG |
| Patchy-F2 | AAGTGCAGTAATTTCCGTGAGG |
| Patchy-F3 | TGCAGTAATTTCCGTGAGGGAA |
| Patchy-F4 | AGTAATTTCCGTGAGGGAATCC |
| Patchy-F5 | AATTTCCGTGAGGGAATCCT |
| Patchy-F6 | TTCCGTGAGGGAATCCTCG |
| Patchy-F7 | CGTGAGGGAATCCTCGAG |
| Patchy-F8 | GAGGGAATCCTCGAGAAGC |
| Patchy-F9 | GGAATCCTCGAGAAGCTTTTC |
| Patchy-F10 | ATCCTCGAGAAGCTTTTCATATTCT |
| Patchy-F11 | CTCGAGAAGCTTTTCATATTCTTTTTGC |
| Patchy-F12 | GAGAAGCTTTTCATATTCTTTTTGCTTG |
| Patchy-F13 | AAGCTTTTCATATTCTTTTTGCTTGGT |
| Patchy-F14 | CTTTTCATATTCTTTTTGCTTGGTGATATC |
| Patchy-F15 | TTCATATTCTTTTTGCTTGGTGATATCAT |
| Patchy-F16 | ATATTCTTTTTGCTTGGTGATATCATTCT |
| Patchy-F17 | TTCTTTTTGCTTGGTGATATCATTCTG |
| Patchy-F18 | TTTTTGCTTGGTGATATCATTCTGAAT |
| Patchy-F19 | TTGCTTGGTGATATCATTCTGAATTCC |

| | |
|---|---|
| Patchy-F20 | CTTGGTGATATCATTCTGAATTCCG |
| Patchy-F21 | GGTGATATCATTCTGAATTCCGAC |
| Patchy-R1 | GAAATGCAGCACAACATCGAC |
| Patchy-R2 | ATGCAGCACAACATCGAC |
| Patchy-R3 | CAGCACAACATCGACCAGG |
| Patchy-R4 | CACAACATCGACCAGGCC |
| Patchy-R5 | AACATCGACCAGGCCAC |
| Patchy-R6 | ATCGACCAGGCCACC |
| Patchy-R7 | GACCAGGCCACCGAG |
| Patchy-R8 | CAGGCCACCGAGGAC |
| Patchy-R9 | GCCACCGAGGACGTAC |
| Patchy-R10 | ACCGAGGACGTACGG |
| Patchy-R11 | GAGGACGTACGGCAGAAC |
| Patchy-R12 | GACGTACGGCAGAACCT |
| Patchy-R13 | GTACGGCAGAACCTGGAAAC |
| Patchy-R14 | CGGCAGAACCTGGAAACC |
| Patchy-R15 | CAGAACCTGGAAACCATCGAG |
| Patchy-R16 | AACCTGGAAACCATCGAGAT |
| Patchy-R17 | CTGGAAACCATCGAGATCCAG |
| Patchy-R18 | GAAACCATCGAGATCCAGAACAT |
| Patchy-R19 | ACCATCGAGATCCAGAACATC |
| Patchy-R20 | ATCGAGATCCAGAACATCGAG |
| Patchy-R21 | GAGATCCAGAACATCGAGCTG |
| Patchy-R22 | ATCCAGAACATCGAGCTGG |
| Patchy-R23 | CAGAACATCGAGCTGGACC |
| Patchy-R24 | AACATCGAGCTGGACCTG |
| Patchy-R25 | ATCGAGCTGGACCTGGC |
| Patchy-R26 | GAGCTGGACCTGGCG |
| Patchy-R27 | CTGGACCTGGCGCG |
| Patchy-R28 | GACCTGGCGCGCAA |
| Patchy-R29 | CTGGCGCGCAAGGA |
| Patchy-R30 | GCGCGCAAGGAGGC |
| Patchy-R31 | CGCAAGGAGGCCCTG |
| Patchy-R32 | AAGGAGGCCCTGGAGG |
| Patchy-R33 | GAGGCCCTGGAGGCG |
| Patchy-R34 | GCCCTGGAGGCGAGC |
| Patchy-R35 | CTGGAGGCGAGCAGG |
| Patchy-R36 | GAGGCGAGCAGGATCAAG |
| Patchy-R37 | GCGAGCAGGATCAAGTCC |
| Patchy-R38 | AGCAGGATCAAGTCCGAG |
| Patchy-R39 | AGGATCAAGTCCGAGTTCCT |
| Patchy-R40 | ATCAAGTCCGAGTTCCTCG |
| Construction of RsmY reporter | |

| | |
|---|---|
| rsmY-F | ATGCAGCAGGCCTCTCGAGGGTACCATCAGGTAGTAGAAGGCGTGC |
| rsmY-R | CAGACCTCTATCCTGACATC |
| PJN105-F | GGGCCCAAGCTTGCTAGCGAATTCCTGCAGCC |
| PJN105-R | GGTACCCTCGAGAGGCCT |
| RNAseIII-mScarlet-F | GATGTCAGGATAGAGGTCTGGGATCCTAACTAACTAGCGATC |
| RNAseIII-mScarlet-R | CTCGCCCTTGCTCACCATAGCTGTTTCCTGTGTGATAAAG |
| mScarlet-F | ATGGTGAGCAAGGGCGAG |
| T1-mScarlet-R | CCTAGGACTGAGCTAGCTGTCAAATCCCCAATTCGATCGTCCG |
| sfGFP-F | ATGCGTAAAGGCGAAGAGCT |
| T1-sfGFP-R | ATTCGCTAGCAAGCTTGGGCCCGCTCCTAGCGGCGGATTTGT |
| J23102-sfGFP-F | TTGACAGCTAGCTCAGTCCTAGGTACTGTGCTAGCTACTAGAGAAAGAGGAGAAATACTAGATGCGTAAAGGCGAAGAGC |
| DsRed-F | TAGGTACTGTGCTAGCTACTAGAGAAAGAGGAGAAATACTAGATGGCCTCCTCCGAGGA |
| DsRed-R | CTACAGGAACAGGTGGTGGC |
| Construction of ClpV1 reporter | |
| clpV1-F | CAGAGATGCGTAATCTTCAAACACTACTACACCGGCCACCTC |
| clpV1-R | CTGCCTCGCCCTTGCTCACCATGCGCAGCTTGCAGAATACC |
| Reporter-F2 | ATGGTGAGCAAGGGCGA |
| Reporter-R2 | GTTTGAAGATTACGCATCTCTGC |
| Construction of HcnB reporter | |
| hcnB-F | CGTAATCTTCAAGAGGTGATGAGCGGTGAACT |
| hcnB-R | CTTGCTCACCATGAAGAGGACGCAGGGGAC |
| Reporter-F3 | TGCGTCCTCTTCATGGTGAGCAAGGGCGA |
| Reporter-R3 | GCTCATCACCTCTTGAAGATTACGCATCTCTGC |
